# Voltage-sensitive dye delivery through the blood brain barrier using adenosine receptor agonist Regadenoson

**DOI:** 10.1101/316505

**Authors:** Rebecca W. Pak, Jeeun Kang, Heather Valentine, Leslie M. Loew, Daniel L.J. Thorek, Emad M. Boctor, Dean F. Wong, Jin U. Kang

**Affiliations:** Biomedical Engineering, Johns Hopkins University School of Medicine, Baltimore, MD, USA; Radiology and Radiological Sciences, Johns Hopkins University School of Medicine, Baltimore, MD, USA; R.D. Berlin Center for Cell Analysis and Modeling, University of Connecticut School of Medicine, Farmington, CT, USA; Electrical and Computer Engineering, Johns Hopkins University | Whiting School of Engineering | Baltimore, MD

## Abstract

Optical imaging of brain activity has mostly employed genetically manipulated mice, which cannot be translated to clinical human usage. Observation of brain activity directly is challenging due to difficulty in delivering dyes and other agents through the blood brain barrier (BBB). Using fluorescence imaging, we have demonstrated the feasibility of delivering the near-infrared voltage-sensitive dye (VSD) IR-780 perchlorate to the brain tissue through pharmacological techniques, via an adenosine agonist (Regadenoson). Comparison of VSD fluorescence of mouse brains without and with Regadenoson showed significantly increased residence time of the fluorescence signal in the latter case, indicative of VSD diffusion into the brain tissue. Dose and timing of Regadenoson were varied to optimize BBB permeability for VSD delivery.

## Introduction

It is established that cellular activity in the brain forms the basis for animal behavior. Yet the complexity of neuronal architecture and the privileged anatomical location of the brain represent long standing impediments to understanding the brain and how best to treat neurological disorders. Current non-invasive techniques rely on bulk effects and thus, do not give cellular resolution. There have been a number of optical imaging approaches to directly monitor brain activity with high spatial and temporal resolution. These include one-photon fluorescence microendoscopy, two-photon confocal microscopy, and endoscopic fiber imaging [1,2]. To visualize brain activity without genetic manipulation, functional fluorescent dyes have been employed, with one of the most widely used types being voltage-sensitive dyes (VSDs) [3]. While VSDs enable individual neuron monitoring with fast time responses and have the ability to operate without modification of the subject, drawbacks include potential toxicity and lower signal-to-noise ratio compared to genetically encoded indicators [4]. Many common VSDs also operate within the visible wavelength range, leading to shallow imaging depth due to absorption. Another challenging aspect of using VSDs is the delivery of dye to the brain.

A variety of methods have been investigated for drug and dye delivery through the blood brain barrier (BBB), a highly-selective semipermeable membrane separating the blood from the brain tissue and extracellular fluid. Osmotic opening of the BBB in which a solution of arabinose or mannitol is infused into the brain has been used for therapeutic applications, but is invasive and causes transient brain edema and gradual dehydration that likely affects neurological activity [5]. Nanocages and nanoparticles have also been considered, but pose potential toxicity due to their persistence in the body [6]. Additionally, many factors influence the release of contents from nanocages and the mechanisms depend on diffusion, osmosis, and degradation that must be tuned [7]. Without proper delivery of the VSD to the brain tissue, there would be no VSD fluorescence reporting neural activity. Focused-ultrasound (FUS) was also considered, but often requires incorporation of the drugs into nanoparticles or microbubbles [8]. There is also some evidence that FUS is accompanied by neuromodulation [9] and since it creates mechanical stress on the cell junctions, risks include hemorrhage and neuron damage [10]. Alternatively, studies have shown that activation of adenosine receptors increases BBB permeability [11]. Thus, to address the delivery obstacle while also facilitating deeper brain imaging, we evaluated the effectiveness as well as the optimal timing and dose of the FDA-approved adenosine receptor agonist Regadenoson (tradename Lexiscan) injected with near-infrared (NIR) VSD IR-780. With the increased BBB permeability due to Regadenoson, the VSD was able to diffuse into the brain tissue, facilitating the monitoring of neuronal cell activity minimally invasively through the intact skull.

## Methods

In this study, IR-780, an NIR cyanine VSD was used to study the effectiveness of Regadenoson. The use of an NIR VSD increased imaging depth and provided the opportunity to image the brain through the intact skull. Due to the delocalized positive charge of the dye, hyperpolarization led to uptake of the dye into the neurons, whereas depolarization resulted in dye leaving the cells [12]. Cyanine dye molecules form non-fluorescent aggregates when in high concentrations, thus leading to quenched fluorescence at the resting membrane potential [13]. On the other hand, action potential firing releases the dye into the extracellular space to cause an increase in the fluorescence signal. Thus, fluorescence imaging of VSDs provides an effective method for monitoring neuron activity. However, VSDs must be delivered to the brain, which can be challenging due to the brain’s immune privilege and BBB.

To test the effects of Regadenoson (Astellas Pharma US, Inc.) on IR-780 perchlorate (576409, Sigma-Aldrich Corp., OS) VSD diffusion, 35-50 g CD1 mice (Charles Rivers Laboratory, Inc., MA, N = 7) were anesthetized with a 50:1 solution of ketamine:xylazine. 24G × ¾” intravenous (iv) tail vein catheters (Terumo Corporation) were placed for injection consistency. To demonstrate the potential for non-invasive procedures, only the scalp skin was removed for the first 3 mice. These mice received iv tail vein injections according to the through-skull fluorescence imaging validation protocol in Fig. 1a.

**Fig. 1.**
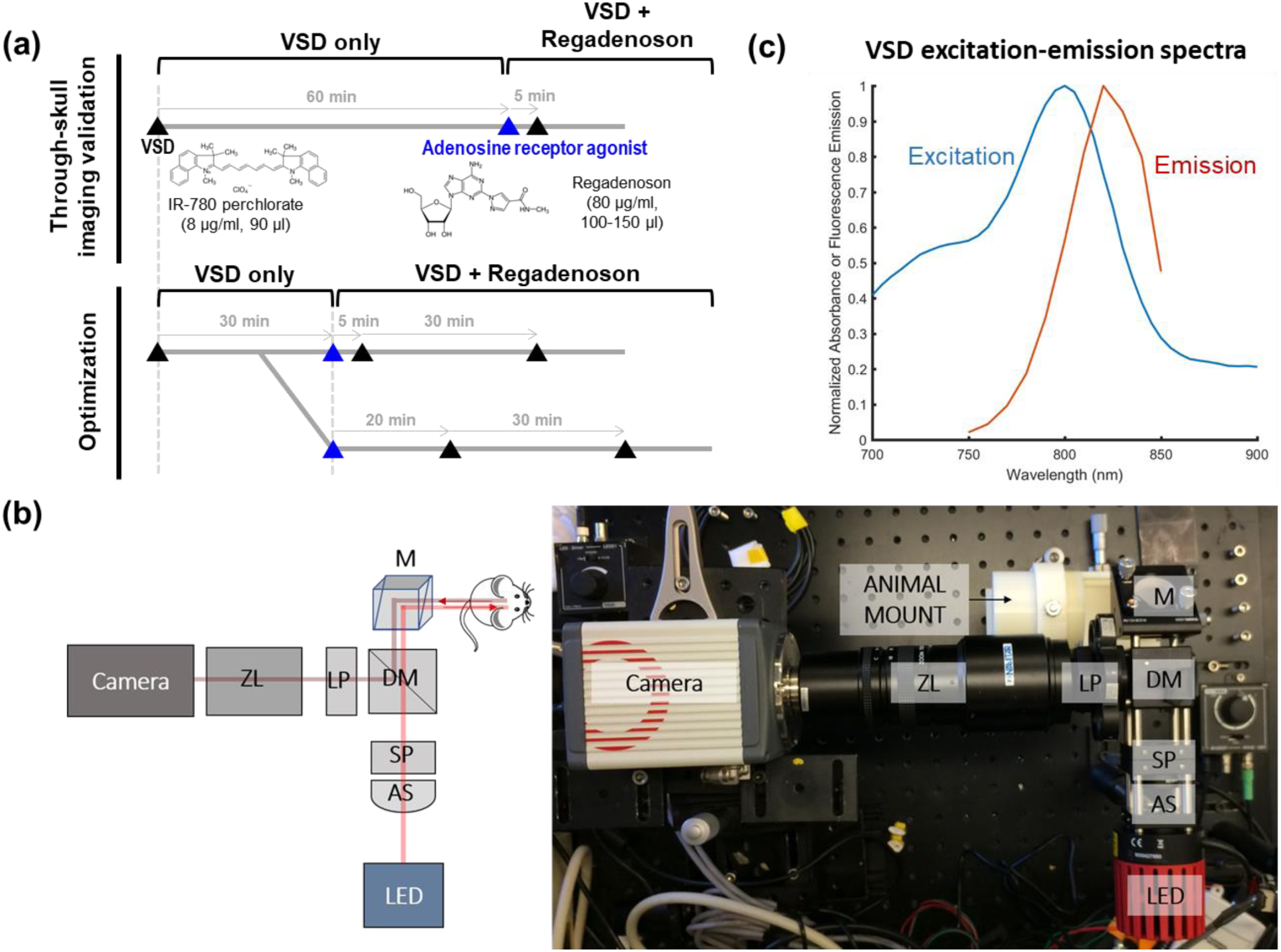
(a) *In vivo* experimental protocol, (b) NIR fluorescence system setup: AS = aspheric lens, SP = shortpass filter, M = mirror, DM = dichroic mirror, LP = longpass filter, ZL = zoom lens, and (c) VSD spectral characteristics.

To verify and quantify dye diffusion, a craniotomy was performed on each of the remaining mice so scattering from the skull could be distinguished from diffusion into the tissue. A head fixation mount was fastened to the skull using dental cement. A dental drill was used to create 6 × 7 mm^2^ cranial windows. The skull piece was removed and replaced by a #1 thick (130-160 μm) glass coverslip cut to size, also secured using dental cement. After recovering from the craniotomy procedures for at least a week, the mice received tail vein injections in a similar schedule as before. However, the time between injections was shortened since the first study with intact skulls showed the fluorescence intensity reaching a steady state around 15-20 min after dye injection. The craniotomy study injections followed the optimization protocol in Fig. 1a.

NIR fluorescence imaging was used to assess dye diffusion, particularly to evaluate optimal injection time and dose of Regadenoson for best dye diffusion into the brain tissue. A typical fluorescence imaging setup (Fig. 1b) was employed with an 800 mW LED light source, operating at a center wavelength of 780 nm and an achromatic lens as a condenser. A shortpass dichroic mirror at 805 nm separated the excitation and emission light. Additional filtration was accomplished with excitation and emission filters. This imaging system enables high-sensitivity sensing of VSD fluorescence with excitation-emission spectra characteristics shown in Fig. 1c. Focusing and magnification was achieved with a zoom lens (Navitar Inc.) attached to an sCMOS camera (Hamamatsu Photonics). For the craniotomy study, mice were head-fixed to avoid motion artifacts from breathing.

To account for differences in injection timing, the time sequences of the VSD only and VSD + Regadenoson conditions were synchronized by aligning *injection peaks* (the maximum of the fluorescence signals, defined in Fig. 3 center panel). Before each dye injection, a background image was taken. After the experiment, the corresponding background image was subtracted from each 30 min sequence of images to examine fluorescence contributions due only to that particular dye injection and reject any fluorescence signal remaining from previous injections or autofluorescence. In order to compare the VSD only condition to the VSD + Regadenoson condition, the *condition subtraction* curve, a differential fluorescence intensity trace between VSD and VSD + Regadenoson conditions (defined in Fig. 3 center panel), was calculated pixel-by-pixel. Regions of interest (ROIs) were chosen to reflect large in-focus vasculature versus microvasculatures. Average VSD fluorescence intensity over the ROIs was plotted.

A histological verification study was conducted using three Sprague Dawley rats weighing 280-350 g. The first rat was injected with Regadenoson alone, the second one was injected with IR-780 VSD, and the third was injected with Regadenoson followed 5 min later by IR-780 VSD. All rats were sacrificed 1 hour post injection(s). The whole brains of the rats were immediately harvested and placed in fresh 10% formalin for > 48 hours with gentle agitation using a conical rotator. Cryoprotection processing was done via a series of sucrose gradients (15%, 20%, 30% for 12-24 hours each). Brains were frozen-sectioned at 300 μm thickness. Slides with tissue sections in ProLong Diamond Anti-face mountant were imaged using the LI-COR Odyssey for fluorescence visualization of VSD perfusion.

## Results

Through-skull fluorescence imaging validated the capability of the system and VSD to be used minimally-invasively. In this first study in which only the scalp was removed and the skull remained intact, the fluorescence signal decay was compared between the VSD only and VSD + Regadenoson conditions. An exponential curve was fit following the model 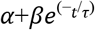 for which the exponential decay coefficient, *γ*, was defined as *γ*=1/τ. As shown in Fig. 2a where the y-axis represents the mean fluorescence signal over the whole mouse brain after normalization, a much faster decay in fluorescence signal was observed without Regadenoson. The faster decay of the fluorescence normalized mean intensity without Regadenoson implies that after the initial peak in fluorescence due to the systemic VSD injection, the VSD remains mostly within the circulation and vessels then decays as the body gets rid of the dye as waste. It should be noted that since IR-780 perchlorate is a small molecule VSD, it is capable of some natural diffusion through the vessels and permeation into the brain tissue as shown by the long background signal. The longer mean time constant when Regadenoson was injected (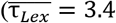 min compared to 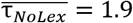 min) suggests that the larger percentage of VSD is penetrating into the brain tissue through the BBB. Hence, the fluorescence signal remained stronger for longer since the dye stuck in the brain tissue exhibited a longer diffusion time of tens of minutes. Thus, the slowed VSD fluorescence decay with Regadenoson suggests a greater BBB penetration. The difference in exponential decay coefficients was compared by the t-test and statistical significance was validated: *p =* 0.0146. Additionally, this trend can be observed in the background subtracted images from each condition at 5-min intervals (Fig. 2b).

**Fig. 2.**
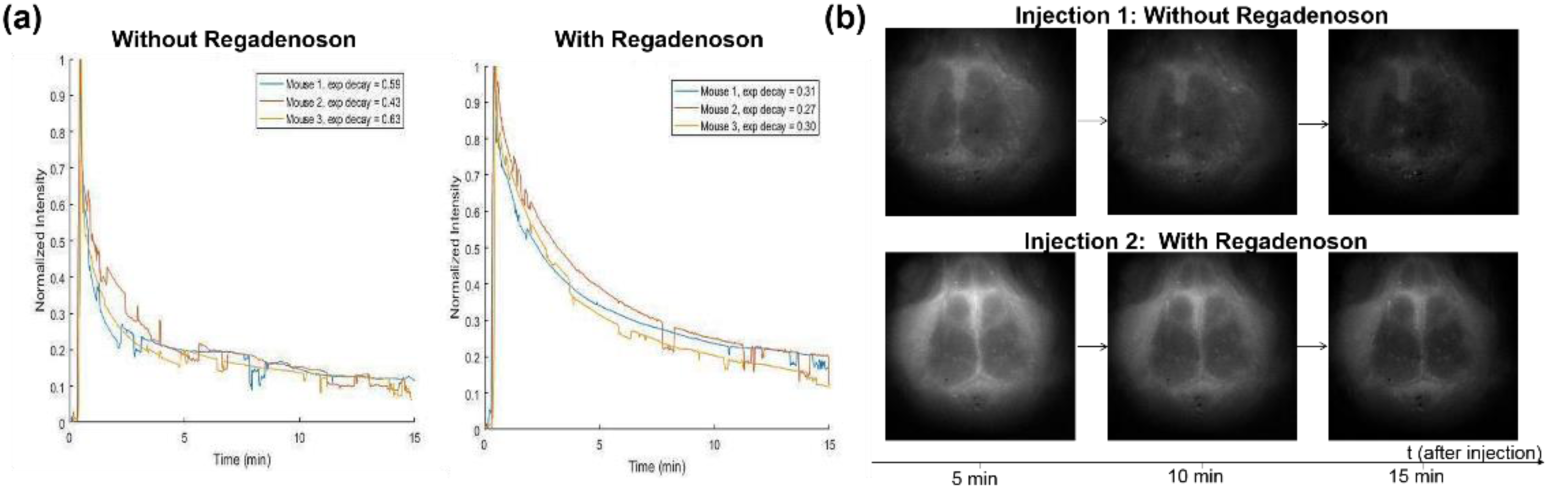
Through-skull (a) decay curves and (b) image time sequence showing fluorescence signal without and with Regadenoson.

**Fig. 3.**
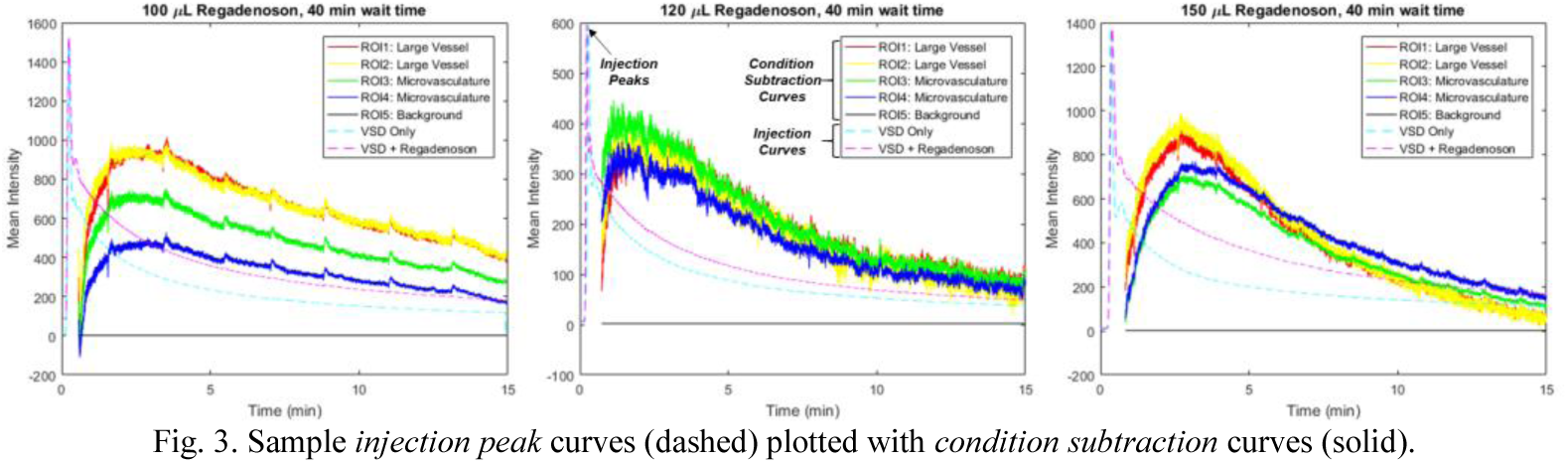
Sample *injection peak* curves (dashed) plotted with *condition subtraction* curves (solid).

Optimization of the dosage and timing of Regadenoson administration was also conducted to achieve optimal VSD diffusion into the brain tissue. After processing and aligning images as outlined in the Methods, the mean fluorescence intensity for *condition subtraction* curves for chosen regions of interest (ROIs) corresponding to large vessels (red and yellow curves), brain tissue and microvasculatures (green and blue), and background (black) were plotted (Fig. 3). The scaled *injection curves* (dashed blue and magenta curves for VSD only and VSD + Regadenoson, respectively) were also plotted to observe the timing. A time delay can be observed between the increase in the *condition subtraction* curves and the *injection peaks*. To define the optimal Regadenoson dose for BBB opening, each VSD fluorescence profile using different Regadenoson doses (i.e., 100, 120, and 150 μl) were fit to an exponential of the form 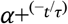 Fig. 4a shows the raw VSD fluorescence profile (right panel) and the exponential curve fitting of differential fluorescence intensity (left panel) obtained at 150 μL Regadenoson dose. In the VSD fluorescence profile, the blue curve indicates the trace with VSD only and the red curve is with VSD + Regadenoson administration. The dashed black vertical lines indicate the range of indices chosen for the proposed curve fitting shown in the left panels at their respective doses. It can be observed that the injection curves generally have 2-3 peaks, so the starting index was found by approximating the full-width-half-max (FWHM) of the first peak and multiplying it by four to account for approximately the end of the second peak. The last index was located about 15 sec after the *condition subtraction* curve reached a maximum value. To capture the differences in VSD perfusion with and without Regadenoson administration, the time constant τ was compared for different Regadenoson doses (Fig. 3 shows the *condition subtraction* curves for each Regadenoson dose with a 40 min wait time. The steepest slope appears in the case of 120 μL of Regadenoson.). The steeper slope or smaller τ implies greater BBB permeability due to faster diffusion and accumulation of the dye in the brain tissue. This contributes to a quicker increase in fluorescence intensity in the brain tissue region and thus a longer residence time and slower decay in terms of overall fluorescence intensity. The range of time constants for the *condition subtraction* curves was found by fixing α and β while varying τ within a tolerance of 2 times the best fit’s root-mean-square error (RMSE). Fig. 4b shows the raw data with 150 μl Regadenoson in blue, the best fit as a solid green curve, and the τ tolerance range which was from −3.4 sec to 3.36 sec as presented in dotted magenta and red curves.

**Fig. 4.**
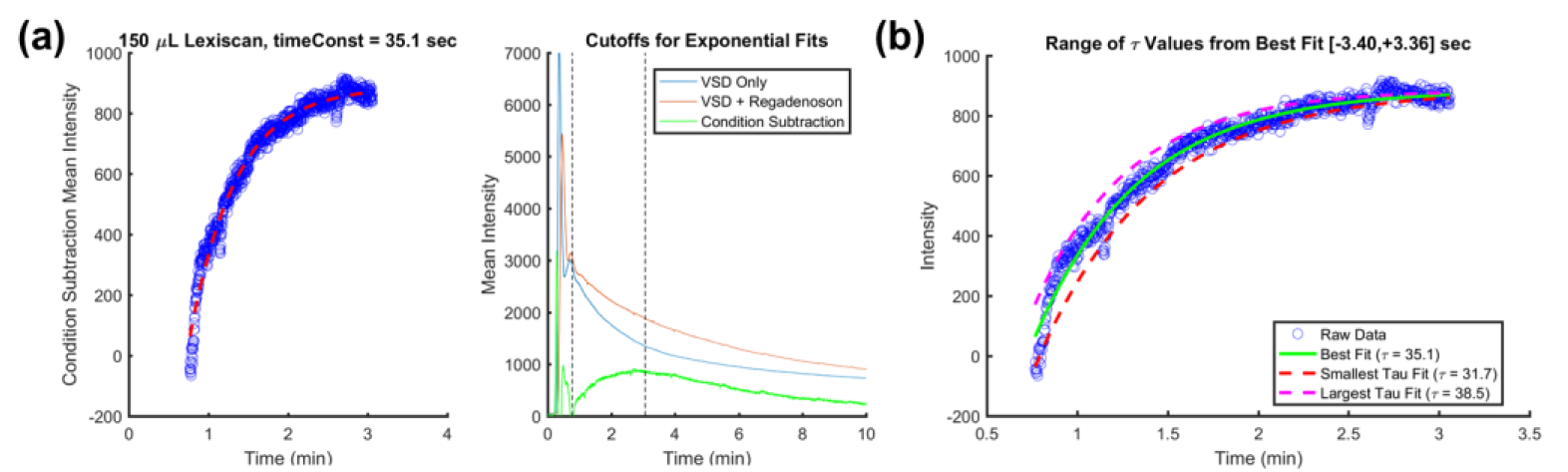
Sample a) exponential curve fitting of the *condition subtraction* curve with 150 μL Regadenoson dose and b) the range of τ values for fits within a tolerance of 2 × RMSE_Bestfit_.

The efficiency of different “wait times” between Regadenoson and VSD injections was also evaluated. When comparing 5 or 20 min wait times to 40 or 55 min wait times (Fig. 5, left), the slope of the initial increase is much steeper in the latter cases. This is also demonstrated by the time constants (τ_5min_ = 29.5 sec, τ_20min_ = 39.2 sec, τ_40min_ = 15.9 sec, τ_50min_ = 5.7 sec), which are inversely proportional to BBB permeability. The result suggests that Regadenoson takes at least a few minutes to start having an effect on BBB permeability and the optimal wait time is 40 min or more.

**Fig. 5.**
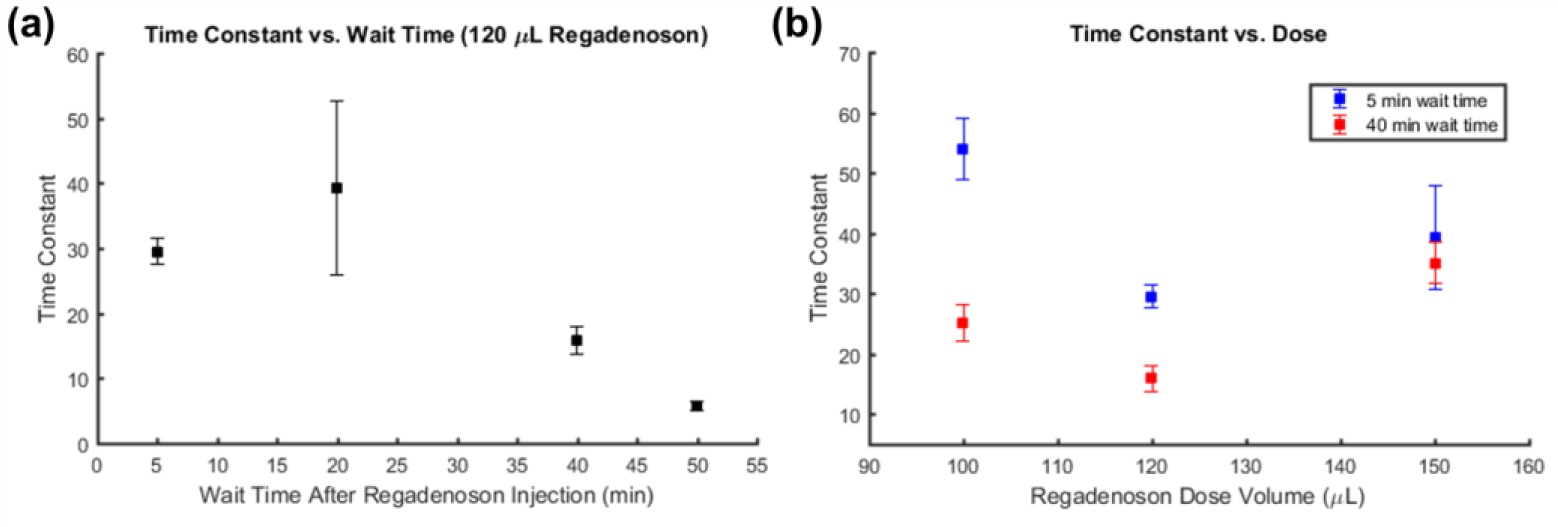
Time constant variation with wait time between injections (left) and with Regadenoson dose (right).

Different doses of Regadenoson were studied by changing the volume of Regadenoson administered: 100, 120, and 150 μL at 5 and 40 min wait times (Fig. 5, right). To minimize side effects from different volumes, a constant total injection volume of 150 μL was maintained by filling any remaining volume with saline. As before, the slope of the *condition subtraction* curves was observed. The highest permeability and optimal dose appeared to occur when using 120 μL of Regadenoson and 30 μL of saline. Interestingly, the wait time between Regadenoson and VSD injections seems to have a diminished effect with increased dose.

A histopathological verification study was conducted to further support the findings. Three rats were injected with Regadenoson only, both Regadenoson and VSD, and VSD only, respectively. Fig. 6 shows the low signal for the first rat, where only a low-level autofluorescence background can be observed. The brain slices of the third rat show substantial VSD fluorescence, but this signal is markedly less compared to the second rat with VSD + Regadenoson. This confirms Regadenoson’s potential as a VSD delivery agent, indicating its ability to increase VSD concentration in superficial and deep brain tissue layers through the BBB to facilitate increased-depth brain imaging.

**Fig. 6.**
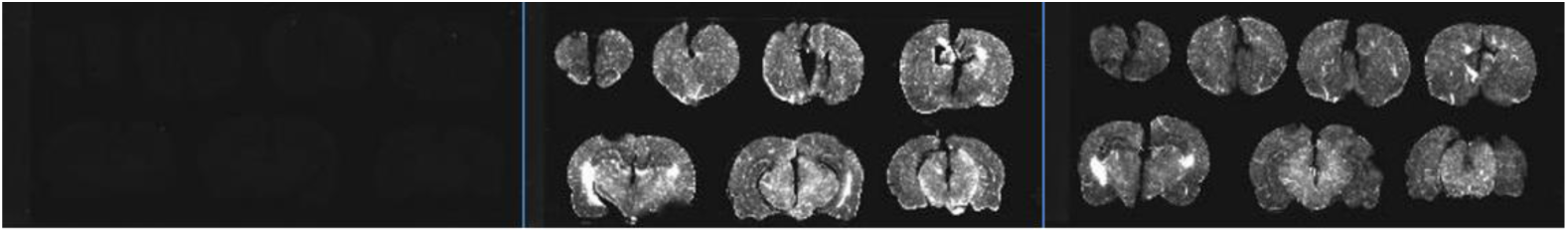
Brain slices with Regadenoson only (left), Regadenoson + VSD (middle), and VSD only (right).

## Discussion and Conclusion

We have presented the effectiveness of adenosine receptor agonist, Regadenoson, in increasing the BBB permeability to allow greater VSD diffusion into the brain tissue for brain activity monitoring applications at increased imaging depths. We have shown that a Regadenoson dose volume of about 120 μL with a wait time of at least 40 min increases the BBB permeability the most.

In our first through-skull fluorescence imaging validation studies with intact skulls, the lack of stereotaxic fixation increased noise in our signal. This is due to the beam not being perfectly uniform in intensity and the fluorescence signal intensity’s dependence on the incident intensity. To control for these movement-induced changes, all the following studies included stereotaxic fixation.

After the first studies, in which the VSD fluorescence was monitored for 1 hour after each dye injection, it was found that the fluorescence intensity reached a steady state before 30 minutes. In order to keep the mouse anatomy constant, the same mouse was used for both conditions (VSD only and VSD + Regadenoson). To account for remaining fluorescence after any previous VSD injections, a background image was taken after 30 min of monitoring when it is assumed that any further decay in fluorescence signal due to previous injections was negligible. This background image was subtracted from images collected from the proceeding VSD injection.

For the optimization of Regadenoson dose and timing studies, the cranial windows varied slightly in their locations for each mouse. However, roughly equivalent ROIs were chosen based on in-focus major vasculature versus out-of-focus microvasculatures. It can also be seen in the results (Fig. 3) that all the ROIs have approximately the same time course trend but varied slightly in their intensities. Equivalent ROIs were also used to measure time constants through exponential curve fitting.

Regadenoson is routinely used for cardiac stress testing, raising a concern about how faster heart rate and circulation affects the results. However, with increased circulation, it would be reasonable to expect faster flushing out of the VSD from the circulatory system. Astellas, the manufacturer of Regadenoson, also warns of changing blood pressure (increase or decrease), which could contribute to the difference in slopes of the *condition subtraction* curves.

Another concern is with regard to dosage based on weight. Clinically, Regadenoson is given to humans at the same concentration and volume regardless of weight or size. Thus, in this study, the same concentration and volume were given regardless of animal weight or size, mimicking the human protocol. We approximated the mouse dose based on the clinical human dose and the ratio between average mouse and human weights. We found this dose was insufficient, possibly due to the higher metabolism, heart rate, etc. of mice as compared to humans. The rat dosage used for histopathological studies that was effective in increasing BBB permeability was similar to that used in the mice, although the rats weighed significantly more and their metabolism is much slower than a mouse. Alternatively, anesthesia may have cardiac implications, which could alter Regadenoson’s dilatory effects. Perhaps rodent dose can be generalized, but for animals with weights more than an order of magnitude different, slight adjustments may need to be made for the optimal conditions.

Overall, this study validated the use of venous injections of Regadenoson for BBB opening to deliver NIR VSD, IR-780 perchlorate. It also demonstrated the possibility of using IR-780 VSD for minimally invasive through-skull fluorescence imaging of neuronal activity. General dosage and timing guidelines were created for optimal VSD diffusion results.

## Funding

Brain Initiative PHS Grant 1 R24 MH106083.

## Disclosures

The authors declare there are no conflicts of interest related to this article.

